# IonQuant enables accurate and sensitive label-free quantification with FDR-controlled match-between-runs

**DOI:** 10.1101/2020.11.02.365437

**Authors:** Fengchao Yu, Sarah E. Haynes, Alexey I. Nesvizhskii

## Abstract

Missing values weaken the power of label-free quantitative proteomic experiments to uncover true quantitative differences between biological samples or experimental conditions. Match-between-runs (MBR) has become a common approach to mitigate the missing value problem, where peptides identified by tandem mass spectra in one run are transferred to another by inference based on m/z, charge state, retention time, and ion mobility when applicable. Though tolerances are used to ensure such transferred identifications are reasonably located and meet certain quality thresholds, little work has been done to evaluate the statistical confidence of MBR. Here, we present a mixture model-based approach to estimate the false discovery rate (FDR) of peptide and protein identification transfer, which we implement in the label-free quantification tool IonQuant. Using several benchmarking datasets generated on both Orbitrap and timsTOF mass spectrometers, we demonstrate superior performance of IonQuant with FDR-controlled MBR compared to MaxQuant (19-38 times faster; 6-18% more proteins quantified and with comparable or better accuracy). We further illustrate the performance of IonQuant, and highlight the need for FDR-controlled MBR, in two single-cell proteomics experiments, including one acquired with the help of high-field asymmetric ion mobility spectrometry (FAIMS) separation. Fully integrated in FragPipe computational environment, IonQuant with FDR-controlled MBR enables fast and accurate peptide and protein quantification in label-free proteomics experiments.

## Introduction

Due to its sensitive and high-throughput nature, liquid chromatography-mass spectrometry (LC-MS) is a popular technology to identify and quantify peptides and proteins from complex samples. Various approaches to LC-MS data acquisition (1–4) have been developed, among which data-dependent acquisition (DDA) remains the most commonly used strategy. In the course of a DDA run, eluted peptides are introduced into a mass spectrometer, where peptide ions are sampled for fragmentation and identified from the resulting tandem mass (MS/MS) spectra. Precursor peptide ion intensities are assumed to be correlated with the actual peptide amount, yielding relative peptide and, after an additional peptide to protein roll-up step, protein quantification. Peptide ions successfully targeted and identified by MS/MS are used to calculate peptide and then protein abundances. However, due to the stochastic nature of intensity-based sampling of peptide ions for MS/MS analysis, not all peptides are consistently identified in all runs. This in turn gives rise to missing quantification values, weakening essential comparisons between different biological samples or experimental conditions. Missing values are generally more prevalent in DDA proteomics than in genomics or transcriptomics. The issue of missing data can be alleviated to some degree using the data-independent acquisition (DIA) strategy (5–9). However, as label-free quantification using DDA data remains popular, there is a critical need to improve computational solutions for this method.

To address the missing value problem in DDA-based proteomics, a number of “identification transfer” approaches have been devised (10–13), exemplified by the match-between-runs (MBR) option in MaxQuant (14, 15) that allows “transfer” of identified precursor peptide peaks from one run (referred to below as donor run) to another (acceptor). Given a peak identified by MS/MS in the donor run, attributes such as m/z, charge state, and retention time, are used to locate a corresponding peak in the acceptor run that is most likely the same peptide. The intensity of the donor peak is then assigned to the acceptor peak, thus filling in the missing value. With more quantified features in common between runs, a greater number of peptides and proteins can be compared among different runs and experiments, increasing the depth of experimental findings (16, 17).

While the goal of MBR is to mitigate the missing value problem, it has the potential to introduce false positives, as transferred peaks have not been rigorously identified using MS/MS spectra in the acceptor run. Lim et al. (18) evaluated the false transfer rate of MBR using a two-organism dataset. They concluded that there was a considerable proportion of false positives from MBR when using MaxQuant, yet most were removed with additional filtering as part of the LFQ calculations. However, in practical settings, even with the additional filtering, FDR of MBR may still be unacceptably high. Thus, this subject deserves a more rigorous treatment that can be generalized across different samples and experimental designs. Here, we propose a semi-supervised approach to control the FDR of MBR, extending our earlier work on FDR for protein identification (4, 19) and DIA quantification (20, 21). We implement FDR-controlled MBR in IonQuant (22), which has been extended to support LC-MS data both with and without ion mobility. We also implement a new protein abundance calculation module in IonQuant based on the MaxLFQ strategy (15), improving upon our previously described top-N approach (21, 22). Using the dataset from Lim et al. (18), we reproduce the authors findings and demonstrate that IonQuant with FDR-controlled MBR has a lower false positive rate and higher sensitivity compared to MaxQuant. With two additional datasets from timsTOF Pro mass spectrometers, we demonstrate that FDR-controlled MBR results in higher quantification precision (lower CV), accuracy, and sensitivity. Finally, we demonstrate that IonQuant displays high sensitivity and precision in single-cell data with or without high-field asymmetric ion mobility spectrometry (FAIMS) separation, and that FDR control for MBR is crucial in such datasets. Overall, we propose an efficient approach to perform MBR with FDR control while maintaining high quantification accuracy and precision. We implement the new methods as a default option in IonQuant, readily available as a standalone tool or within our integrated computational platform FragPipe (https://fragpipe.nesvilab.org/).

## Experimental Procedures

### Experimental Design and Statistical Rationale

We used five datasets in this work. In all datasets, we estimated the identification false-discovery rate using the target-decoy approach (4). For MSFragger, PSMs, peptides, and proteins were filtered at 1% PSM and 1% protein identification FDR. For MaxQuant, PSMs and peptides were filtered at 1% PSM FDR, and proteins were filtered at 1% protein FDR, which is MaxQuant’s default setting. A two-organism dataset (*H. sapiens* and *S. cerevisiae*) with 40 LC-MS runs from Lim et al. (18) was generated on an Orbitrap Fusion Lumos mass spectrometer (Thermo Fisher Scientific). In this dataset, 20 runs include only *H. sapiens* proteins, while the remaining 20 runs contain a mixture of *H. sapiens* and *S. cerevisiae* proteomes. *S. cerevisiae* peptides transferred to the 20 *H. sapiens*-only runs by MBR are false positives and were used to evaluate the false positive rate. We also employed two datasets from timsTOF Pro (Bruker), as in our previous work (22). A HeLa dataset with 4 replicate injections from Meier et al. (23) was used to evaluate the sensitivity (i.e., quantified protein count) and precision (i.e., coefficient of variation (CV)) of quantification across replicate runs. A three-organism timsTOF dataset (*H. sapiens*, *S. cerevisiae*, and *E. coli*) with 6 runs from Prianichnikov et al. (24) was used to evaluate quantification accuracy, and contains two experimental conditions with ground truth protein ratios: 1:1 (*H. sapiens*), 2:1 (*S. cerevisiae*), and 1:4 (*E. coli*). A single-cell dataset published by Williams et al. (25) was generated on an Orbitrap Fusion Lumos mass spectrometer (Thermo Fisher Scientific). This dataset contains 3 replicate runs with 0 cell (blank runs), 11 replicates with 1 cell, 4 replicates with 3 cells, 4 replicates with 10 cells, and 4 replicates with 50 cells. Numbers of quantified peptides and proteins were used to evaluate sensitivity, and quantification CV was used to evaluate precision. The last dataset was also from a HeLa single-cell experiment (26), acquired on an Orbitrap Eclipse Tribrid mass spectrometer with the help of FAIMS separation. There are 3 single HeLa cell runs, 3 blank runs, and 3 library runs generated from 100 cells. Numbers of quantified proteins were used to evaluate sensitivity.

### Indexing-based MBR

We developed a fast MBR algorithm based on indexing. In IonQuant (22), an index of each run is built and written to the disk for fast feature extraction, which supports data with and without ion mobility information. The peak tracing and normalization modules were improved to make it more sensitive and robust compared to the initial release of IonQuant. The new version performs resampling to make the peaks have the same time interval. Then, it performs Savitzky-Golay smoothing (27), finds the boundaries, and subtracts background noise using Skyline’s approach (https://skyline.ms/wiki/home/software/Skyline/page.view?name=tip_peak_calc). In the normalization module, the whole m/z range is now divided into 10 bins with the same number of ions, which makes normalization more robust for sparse data or samples with large differences in abundance.

Given a run with possible missing values that will accept ions (acceptor run) and a separate run that will be used to fill these missing values (donor run), correlations between the two runs are calculated using overlapped ions’ retention times, intensities, and ion mobilities if applicable: (*o* × *r*_1_ + *o* × *r*_2_)/ 2 or (*o* × *r*_1_ + *o* × *r*_2_ + *o* × *r*_3_)/ 3, where *o* is the overlapping ratio (28); *r*_1_, *r*_2_, and *r*_3_ are Spearman’s rank correlation coefficients of retention time, intensity, and ion mobility, respectively. Up to *n* (user-specified ‘MBR top runs’ parameter, 10 by default) donor runs with the highest correlations (which must be greater than user-specified ‘MBR min correlation’ parameter, 0 by default) are selected.

For each ion in every selected donor run, we locate the target region within the acceptor run using an approach similar to FlashLFQ (29). First, pairs of retention times from the corresponding ions are collected and sorted according to the value from the donor run. Using *d*_*i*_ and *a*_*i*_ to denote the retention times of *i*-th pair of ions from the donor and acceptor runs, respectively, we have pairs from (*d*_1_, *a*_1_) to (*d*_*N*_, *a*_*N*_) sorted by *d*_*i*_, where *N* is the number of overlapped ions. Given a donor ion with retention time *t*, we find its position in the sorted pairs satisfying *d*_*i*_ ≤ *t* < *d*_*i*+1_. Then, we collect all pairs satisfying *d*_*i*_ − τ ≤ *d*_*j*_ ≤ *d*_*i*_ + τ, where τ is a predefined tolerance (‘MBR RT window’ parameter, 1 minute by default). With those pairs, we generate a list whose elements are *a*_*j*_ − *d*_*j*_, and calculate the median (*m*) and median absolute deviation (σ) of that list. The possible target range in the retention time dimension is then:

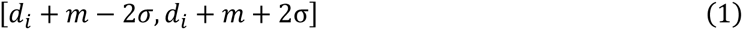

If ion mobility data are used, we take the same approach to locate the target range in the ion mobility dimension (controlled by the ‘MBR IM window’ parameter, 0.05 by default). The transferred ion’s m/z equals the donor ion’s m/z adjusted by mass calibration error (mass calibration is performed by MSFragger (30)). After locating the target region in m/z, retention time, and ion mobility if applicable, we trace all peaks within the region using our recently described algorithm (22). Two isotope peaks (+1 and +2) are also traced to check the charge state and the isotope distribution. Peak boundaries are allowed to extend beyond the target region’s retention time and ion mobility bounds. Peak tracing is performed rapidly using the index, after which the donor ion’s peptide information is assigned to the traced monoisotopic peak.

IonQuant can automatically detect if the data were acquired using FAIMS. If FAIMS was used, IonQuant builds separate spectral indexes corresponding to each compensation voltage. Then, peak tracing, ion detection, and ion transfer are performed within each compensation voltage.

### MBR false discovery rate estimation

To estimate the rate at which false transfers occur, we adopted a supervised semi-parametric mixture model that we previously applied in a number of related applications (19, 20). For each successfully transferred donor ion (i.e., target ion), we try to transfer a decoy ion, created to have the same retention time and ion mobility (if applicable) but with a large m/z shift (31–33). To generate a decoy, we first shift the m/z by +11×1.0005 Th. If there is no traceable peak in that region, we keep decreasing the m/z shift by 1.0005 Th until we successfully trace a peak or until the m/z shift reaches +4 Th.

For all transferred target and decoy ions, we calculate four (without ion mobility) or five (with ion mobility) scores (Table 1). For one of these scores (using the 0/+1/+2 peaks), Kullback-Leibler divergence is used to compare the quality of the traced isotopic distribution to a theoretical one given m/z and charge state, where the Poisson distribution is used as theoretical (34).

**Table 1.** List of individual scores used to compute the composite score for each transferred ion.

| Score | Explanation |
| --- | --- |
| Log10(intensity) | Log-transformed intensity of a traced peak. The intensity can be from an area (without ion mobility) or a volume (with ion mobility). |
| Log10(KL) | Log-transformed Kullback-Leibler divergence of an experimental isotope distribution and the theoretical isotope distribution. 0, +1, and +2 isotope peaks are used. The absolute value is also square root transformed. |
| Abs(ppm) | Absolute value of the mass error (in ppm) from a traced peak. The value is also square root transformed. |
| IM diff | Ion mobility difference between an acceptor ion and its donor ion. The value is also square root transformed. |
| RT diff | Retention time difference between an acceptor ion and its donor ion. The value is also square root transformed. |

We classify all transferred ions (identified with sequence, charge, and modification information) into four types: a target ion that has not been identified by MS/MS in the acceptor run (type 1); a decoy ion that is from a m/z-shifted type 1 ion (type -1); a target ion that has already been identified by MS/MS (type 2); or a decoy ion that is from a m/z-shifted type 2 ion (type -2). Following the strategy we previously used for DIA data (20), we train a linear discriminant analysis (LDA) model using scores from type 2 and -2 ions. From the trained LDA, we calculate a final score for each type 1 and -1 ion:

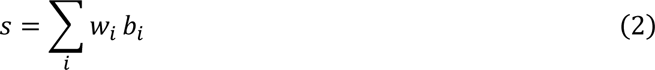

where *s* is the final score, *w*_*i*_ are the weights from LDA, and *b*_*i*_ are the scores detailed in Table 1. If multiple ions were transferred to one location, the top scoring one is kept.

Using the final scores from type 1 and -1 ions, we estimate a posterior probability of correct identification transfer by fitting a mixture model:

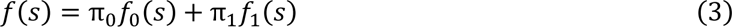

where *f*_0_ is the distribution of correctly transferred ions, *f*_1_ is the distribution of incorrectly transferred ions, π_0_ and π_1_ are the respective priors of false and true transferred ions. We use the expectation-maximization (EM) algorithm (20) to estimate the coefficients and distributions in Equation (3).

After fitting the mixture model, we calculate a posterior probability for each transferred ion using

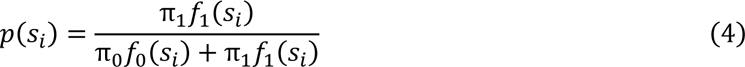

where *s*_*i*_ is the score of the transferred ion. Then, we calculate an ion-level MBR FDR using the posterior probability (35) of type 1 ions:

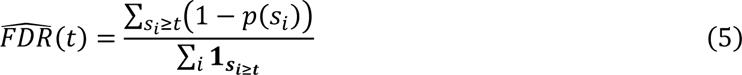

where *t* is a score threshold and ∑_*i*_ **1**_***si***≥***t***_ is the number of type 1 ions whose score is larger than *t*. We can also calculate peptide-and protein-level FDR for MBR by collapsing ions with the same sequence or protein and using the highest probability entry in the FDR calculation.

### Calculating protein intensity using MaxLFQ algorithm

Cox et al. proposed MaxLFQ (15) algorithm to calculate protein intensity with peptide intensities. It has a high precision (low CV) according to our previous study (22). We implemented it in IonQuant to provide a new (default) option in addition to the top-N approach.

Given a study with *N* experiments (samples), and a protein with *M* quantified peptide ions, for each peptide ion *p* ∈ [1, *M*] we calculate a log-ratio of its intensities between experiments *i* and *j*:

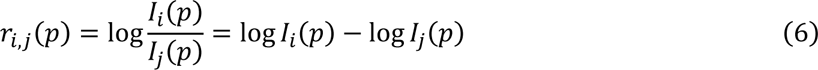

where *I*_*i*_(*p*) is the intensity of peptide ion *p* from *i*-th experiment. If the ion is not quantified in experiment *i* or *j*, we do not calculate the corresponding log ratio. Then, we have a linear relationship among the log-transformed protein intensities and their peptide ion log-ratios:

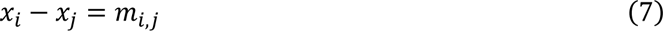

where *x*_*i*_ is the (unknown) log-transformed protein intensity in *i*-th experiment and *m*_*i*,*j*_ is the median of the log-ratios *r*_*i*,*j*_(*p*) among all peptide ions *p* from 1 to *M*. Given the set of 1 to *N* experiments, Equation (7) can be expressed in a matrix form

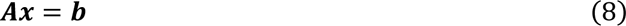

Where

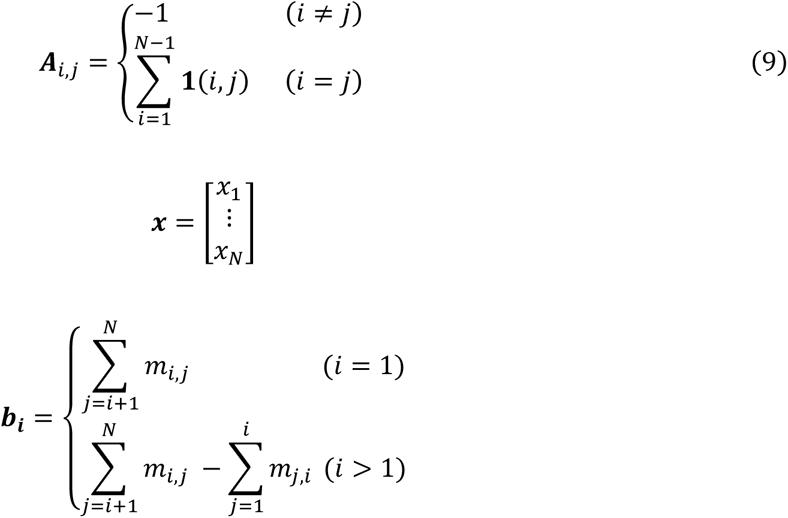

In Equation (9), **1**(*i*, *j*) equals 1 if there is a peptide ion quantified in both experiment *i* and *j*, and 0 otherwise. Equation (8) can be efficiently solved with Cholesky decomposition to get the log-transformed protein intensity *x*_*i*_. Then, the protein intensity in experiment *i* equals *e*^*xi*^.

### Validation of the FDR for MBR approach using two-organism dataset

We used 40 runs from Lim et al. (18) (ProteomeXchange (36) identifier PXD014415) to evaluate the sensitivity and precision of FDR-controlled MBR. This dataset contains 20 runs with only *H. sapiens* proteins and 20 with a mixture of *H. sapiens* (90%) and *S. cerevisiae* (10%) proteins, all acquired on an Orbitrap Fusion Lumos mass spectrometer. Further sample preparation and data acquisition details can be found in the original publication (18). We used FragPipe (version 13.0) with MSFragger (37) (version 3.0), Philosopher (38) (version 3.2.7), and IonQuant (22) (version 1.5.5) to analyze this dataset. For this analysis pipeline, raw spectral files were first converted to mzML using ProteoWizard (version 3.0.20066) with vendor’s peak picking. We used MaxQuant (39) (version 1.6.14.0) and also Skyline (40) (version Skyline-daily (64-bit) 20.2.1.315 (3785d2eb9)) for comparison. We used raw spectral files for MaxQuant and spectral files converted to the mzML format for other tools. A protein sequence database of reviewed *H. sapiens* (UP000005640) and *S. cerevisiae* (UP000002311) from UniProt (41) (reviewed sequences only; downloaded on Jan. 15, 2020) and common contaminant proteins (26448 proteins total) was used. For the MSFragger analysis, precursor and (initial) fragment mass tolerance were set to 50 ppm and 20 ppm, respectively. Reversed protein sequences were appended to the original database as decoys. Mass calibration and parameter optimization were enabled. Isotope error was set to 0/1/2, and one missed trypsin cleavage was allowed. Peptide length was set from 7 to 50, and peptide mass was set to 500 to 5000 Da. Oxidation of methionine and acetylation of protein N-termini were set as variable modifications. Carbamidomethylation of cysteine was set as a fixed modification. Maximum allowed variable modifications per peptide was set to 3. Philosopher (38) with PeptideProphet (42) and ProteinProphet (43) was used to estimate identification FDR. The PSMs were filtered at 1% PSM and 1% protein identification FDR. Quantification and MBR was performed with IonQuant. The minimum number of ions parameter required for quantifying a protein was set to 2 (default). To test the performance of FDR control for MBR, the maximum number of runs used for transfer was set to 40, and the minimum required correlation between the donor and acceptor run was set to 0. Ion-, peptide-, and protein-level MBR FDR thresholds were all set to 1% unless otherwise noted. Protein intensities were computed using the re-implementation of MaxLFQ protein intensity calculation algorithm described above. Default values were used for all the remaining parameters. For MaxQuant comparisons, the parameters were set as close to those described above as possible, with maximum modifications per peptide set to 3, maximum missed cleavages set to 1, LFQ enabled with default settings, maximum peptide mass set to 5000, built-in contaminant proteins were not used, and the second peptide option was not used. Default values were used for all the remaining MaxQuant parameters.

For Skyline comparisons, pep.xml files from PeptideProphet were loaded with probability threshold 0.9486 that corresponds to 1% peptide-ion level FDR in this dataset. A protein FASTA file filtered with 1% protein FDR was also loaded to make sure that Skyline was processing the peptides additionally filtered with 1% protein FDR. Retention time filtering tolerance was set to 0.4 minutes, the same tolerance as in IonQuant. After loading all PSMs, we let Skyline generate decoys by reversing the sequences and shifting the precursor masses. Then, we reintegrated the peaks by training a model with the built-in mProphet (44). Finally, we exported a peptide quantification report with estimated q-values, and filtered the data using 0.01 threshold.

We classified a peptide as an *S. cerevisiae* peptide if it only maps to *S. cerevisiae* proteins. We classified a peptide as *H. sapiens* if it maps to at least one *H. sapiens* protein. The classification was done based on the protein name in the searched protein sequence database: those ending with “_HUMAN” were classified as *H. sapiens* proteins and those ending with “_YEAST” were classified as *S. cerevisiae* proteins.

### Quantification precision comparison using four HeLa cell lysate replicates

We used four replicate HeLa cell lysate runs acquired on a timsTOF Pro mass spectrometer (23) with 100 ms TIMS accumulation time to evaluate quantification precision when MBR is used. As in the previous section, we used FragPipe (version 13.0) with MSFragger (version 3.0), Philosopher (version 3.2.7), and IonQuant (version 1.5.5) to analyze this dataset. MaxQuant (version 1.6.14.0) was used to perform a benchmark comparison. Raw spectral files (.d extension) were used. The sequence database contained reviewed *H. sapiens* (UP000005640) proteins and common contaminants from UniProt (downloaded on Sep. 30, 2019; 20463 sequences). The minimum number of ions parameter required for quantifying a protein was set to 2 unless otherwise noted. For MBR in IonQuant, MBR top runs parameter was set to 3 and MBR min correlation was set to 0. Ion-, peptide-, and protein-level MBR FDR threshold were set to 1%. Remaining parameters were identical to those in the previous section. We used the number of proteins quantified in at least two runs and quantification CV across replicates to evaluate the performance.

### Quantification accuracy comparison using the three-organism dataset

We used the three-organism dataset by Prianichnikov et al. (24) to demonstrate the accuracy of IonQuant with MBR. There are six runs from two experimental conditions (A and B) in which *H. sapiens*, *S. cerevisiae*, and *E. coli* proteins are mixed at known ratios. The ratios between conditions A and B are 1:1 (*H. sapiens*), 2:1 (*S. cerevisiae*), and 1:4 (*E. coli*). These data were acquired on a timsTOF Pro mass spectrometer, and details of the sample preparation and data generation can be found in the original publication (24). We used FragPipe (version 13.0) with MSFragger (version 3.0), Philosopher (version 3.2.7), and IonQuant (version 1.5.5) to analyze the data. MaxQuant results published by Prianichnikov et al. (24) were used as a benchmark comparison. Using the latest MaxQuant (version 1.6.14.0), a reviewed UniProt protein sequence database, and parameters closest to those of MSFragger and IonQuant yielded results similar to those in the original publication (**Supporting Figure S1**). A combined database of reviewed *H. sapiens* (UP000005640), *S. cerevisiae* (UP000002311), and *E. coli* (UP000000625) sequences from UniProt (30788 sequences downloaded Apr. 18, 2020) was used. Ion-, peptide-, and protein-level MBR FDR thresholds were set to 1%. The minimum number of ions parameter required for quantifying a protein was set to 2. Allowed missed cleavages was set to 2, and all other parameters were the same as those in the previous section. We used LFQbench (45) to plot the protein quantification results.

### Single-cell dataset analysis

We used 26 runs published by Williams et al. (25) to demonstrate IonQuant’s performance with single-cell data. There are 3 replicates containing 0 cells which served as negative controls, 11 replicates containing 1 cell, 4 replicates containing 3 cells, 4 replicates containing 10 cells, and 4 replicates containing 50 cells. The data were generated on an Orbitrap Fusion Lumos mass spectrometer (Thermo Fisher Scientific) over a 30 minute LC gradient, with MS/MS spectra acquired in the ion trap. Details of the sample preparation and data acquisition can be found in Williams et al. (25). The raw data files were converted to mzML format using ProteoWizard (version 3.0.19302) with vendor’s peak picking. We used FragPipe (version 13.0) with MSFragger (version 3.0), Philosopher (version 3.2.7), and IonQuant (version 1.5.5) to analyze the data. We also used MaxQuant (version 1.6.14.0) as a benchmark. The database was downloaded along with the data (20129 proteins, ProteomeXchange (36) identifier MSV000085230). In MSFragger analysis, common contaminants and reversed protein sequences were appended by Philosopher. In MaxQuant analysis, the built-in contaminant sequences were used. The precursor mass tolerance was set to 20 ppm, and the initial fragment mass tolerance was set to 0.6 Da. Two missed cleavages were allowed. IonQuant (version 1.5.5) with and without MBR was used. The MBR top runs parameter for MBR transfer was set to 26 and the minimum required correlation was kept at 0. The MaxLFQ protein intensity calculation algorithm was used. The minimum number of ions parameter required for quantifying a protein was set to 1. Multiple ion-level MBR FDR thresholds were applied. The rest of the parameters are the same as those used in the previous section. MaxQuant’s parameters were set as close as possible to those used in MSFragger and IonQuant. We used the numbers of quantified peptides and proteins to evaluate the sensitivity, and we used CV to evaluate the precision of label free quantification with MBR.

### Single-cell FAIMS dataset analysis

We used 9 runs published by Cong et al. (26) to demonstrate the performance of analyzing single-cell data from an Orbitrap Eclipse Tribrid mass spectrometer (Thermo Fisher Scientific) coupled with FAIMS. There are 3 single HeLa cell runs, 3 blank runs served as negative controls, and 3 runs with 100 HeLa cells that served as a library for MBR. Each run has two compensation voltages: -55 V and -70 V. The sequence database contains reviewed *H. sapiens* (UP000005640) proteins and common contaminants from UniProt (downloaded on Sep. 30, 2019; 20463 sequences). We used FragPipe (version 13.0) with MSFragger (version 3.0), Philosopher (version 3.2.7), and IonQuant (version 1.5.5) to analyze the data. Raw spectral files were first converted to the mzML format using ProteoWizard (version 3.0.20253) with vendor’s peak picking. The number of allowed donor runs was set to 9. The rest of the parameters are the same as those used in the previous section. MaxQuant (version 1.6.14.0) was used for comparison. Since MaxQuant does not support FAIMS data natively, we split each raw file into separate mzXML files using FAIMS-MzXML-Generator (https://github.com/PNNL-Comp-Mass-Spec/FAIMS-MzXML-Generator). Scans in each mzXML file have the same compensation voltage (46). Then, we assign fraction number 1 to the mzXML files with compensation voltage equal to -55 V, and fraction number 3 to the mzXML files with compensation voltage equal to -70 V (**Supporting Figure S3**). In this way, ions are only allowed to be transferred among the files with the same compensation voltage. The rest of the parameters were set as close as possible to those used in MSFragger and IonQuant. We compared the number of quantified proteins with and without MBR from MaxQuant and IonQuant.

### Run time comparison

We used the two-organism dataset with 40 Orbitrap Fusion Lumos runs and the HeLa dataset with 4 timsTOF Pro runs to demonstrate the speed of label-free quantification coupled with FDR-controlled MBR in IonQuant (version 1.5.5). MaxQuant (version 1.6.14.0) was used for comparison. For the two-organism dataset, we used a combined database of reviewed *H. sapiens* (UP000005640) and *S. cerevisiae* (UP000002311) sequences from UniProt (41) plus common contaminants (26448 proteins downloaded Jan. 15, 2020). For the HeLa dataset, a database of reviewed *H. sapiens* (UP000005640) proteins from UniProt (20463 proteins downloaded on Sep. 30, 2019) and common contaminants was used. Reversed proteins sequences were appended to both databases as decoys for MSFragger analysis. All other parameters are identical to those used in the previous section. All analyses were run on a desktop with 4 CPU cores (Intel Xeon E5-1620 v3, 3.5 GHz, 8 logical cores) and 128 GB memory. We isolated quantification-specific run times from MaxQuant log files.

## Results and Discussion

### FDR-controlled MBR

We developed an MBR module in IonQuant enabling accurate and fast label-free quantification with match-between-runs peptide ion transfer with the help of the indexing functionality in IonQuant (see Figure 1 for an overview). For each experiment (acceptor run) in the analysis, ion-level Spearman’s rank correlation coefficients with all other experiments are calculated, where an ion is defined as the combination of peptide sequence, modification pattern, and charge state. The percentage of ions overlapping between two runs is used as a weight in the calculation (28). For each acceptor run, IonQuant picks the top N runs with a correlation larger than a certain threshold as donor runs. Both parameters (‘MBR top runs’ and “MBR min correlation’ can be adjusted by the user). Given an ion from a donor run, IonQuant locates a region in the acceptor run where the transferred ion is likely to be using m/z, retention time, and ion mobility (if applicable) distributions from both runs (see Figure 1 and **Experimental Procedures**). For simplicity, we use retention time to describe the region-finding process. Given an ion from a donor run, all ions within a predefined retention time tolerance are collected. Retention time differences from pairs of ions overlapping between the runs are calculated, and the median and median absolute deviation of these differences are found. Then the region for transfer is determined using Equation (1). We use the same approach to locate the ion mobility region. After getting a 1-D (without ion mobility) or 2-D (with ion mobility) region, IonQuant traces peaks using the donor ion’s m/z, taking any mass calibration correction into account. In addition to the monoisotopic peak, two additional isotope peaks (+1 and +2) are also included in peak tracing so that the isotopic distribution and charge state can be used in the evaluation. Finally, IonQuant assigns the donor ion’s peptide to each traced peak and calculates four (without ion mobility) or five (with ion mobility) scores (Table 1) measuring the quality of the peptide ion transfer.

**Figure 1.**
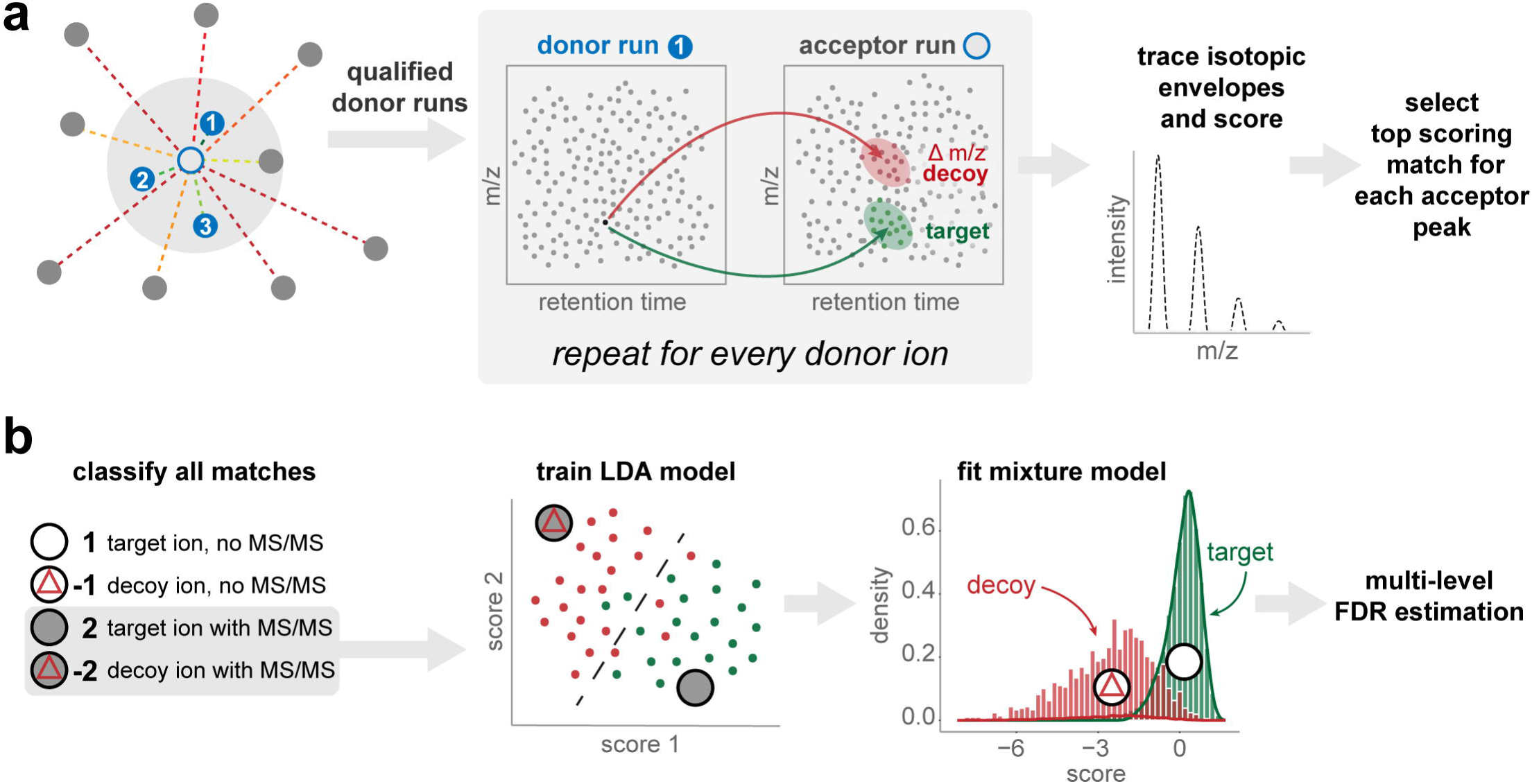
**(a)** Overview of match-between-runs in IonQuant. For each acceptor run (unfilled central point with blue outline) ion-level correlations with all other runs (filled blue and gray points) are calculated, where distance from the central point represents correlation. The top N runs (numbered blue points) within the correlation threshold (gray area) are selected as eligible donor runs. For every ion in each eligible donor run, target and decoy (m/z-shifted) transfer regions are located using retention time (and ion mobility if applicable). Peak tracing in the acceptor run is used to determine the isotopic distribution and the charge state. All matches are evaluated, and the top scoring donor for each acceptor peak is selected for transfer. **(b)** All matches/transferred ions are classified into one of the four categories shown. Type 2 and -2 matches are used to train a linear discriminant analysis (LDA) model. The trained LDA is then used to calculate the final score for type 1 and -1 matches. A posterior probability of correct transfer is estimated by fitting a mixture model, allowing estimation of ion-, peptide-, and protein-level false discovery rate (FDR) for match-between-runs.

In conventional MBR, most notably in MaxQuant, ions matching tolerance criteria are transferred without statistically assessing the confidence in the transfer. Here, we propose a semi-parametric mixture-modeling approach to estimate the FDR of transferred ions (see **Experimental Procedures**). Briefly, decoy ion transfers are generated by transferring ions with an m/z shift. All transferred ions are classified into four types: the ion has not been identified by MS/MS (type 1); the ion is a decoy type 1 ion (type -1); the ion has been identified by MS/MS (type 2); and the ion is a decoy type 2 ion (type -2). IonQuant trains a linear discriminant analysis (LDA) model with type 2 and -2 ions to separate the target and decoy ions. Using the trained model, a final score is calculated for each of the type 1 and -1 ions (Equation (2)). A mixture model (Equation (3)) is built using type 1 and -1 ions, and the expectation-maximization (EM) algorithm is used to fit the model and subsequently calculate the posterior probability. Finally, global ion-level FDR (Equation (5)) is calculated using the local FDR, equal to one minus the posterior probability (Equation (4)). IonQuant also calculates peptide and protein level FDR by collapsing ions with the same peptide and protein, respectively.

In the remainder of the manuscript, we demonstrate the accuracy of FDR-controlled MBR using a two-organism dataset, and the precision and accuracy of subsequent label-free quantification by using HeLa replicate runs, a three-organism dataset, and two single-cell dataset, respectively.

### Evaluation of FDR-controlled MBR method

We used the dataset published by Lim et al. (18) to evaluate the false positive rate of FDR-controlled MBR (see **Experimental Procedures**). The dataset is comprised of 20 LC-MS files from *H. sapiens*-only proteins (“H”) and 20 from a mixture of *H. sapiens* (90%) and *S. cerevisiae* (10%) proteins (“HY”). With MBR, *S. cerevisiae* peptides transferred from HY to H runs are known to be false positives, and can be used to evaluate the false positive rate, equal to false positives (*S. cerevisiae* peptides in H runs) divided by negatives (*S. cerevisiae* peptides in total). To ensure all *S. cerevisiae* peptides in the HY runs have the chance to be transferred, the number of top runs used in transferring was set to 40 and minimum required correlation was set to 0. In evaluation, a peptide was assigned to *S. cerevisiae* if all proteins it maps to are from *S. cerevisiae*, or to *H. sapiens* if at least one of its proteins is from *H. sapiens*.

Overall, IonQuant coupled with MSFragger identified 45875 unique *H. sapiens* peptides and 4610 unique *S. cerevisiae* peptides, ∼19% and ∼31% more *H. sapiens* and *S. cerevisiae* peptides compared to MaxQuant, respectively (Table 2**, Supporting Table S1**). More peptides were also identified or transferred in individual runs with MSFragger and IonQuant. In transferring ions between the runs, IonQuant had a lower false positive rate than MaxQuant, 2.3% compared to 2.7%. The numbers listed for MaxQuant in Table 2 differ slightly from Figure S1 in Lim et al. (18) because of small differences in data analysis settings and version of the tools used. Figure 2 shows average peptide coverage, average peptide false positive rate, average protein coverage, and average protein false positive rate with respect to different MBR FDR thresholds. The peptide/protein coverage values shown are *H. sapiens* peptides/proteins in each H run divided by total *H. sapiens* peptides/proteins identified in the dataset. Peptide coverage increases from 57% to 79% with the inclusion of MBR, and protein coverage increases from 87% to 96%. As the MBR FDR threshold is increased, neither peptide nor protein coverage increase significantly, indicating most *H. sapiens* peptides have been successfully transferred by IonQuant already at 1% MBR FDR. The false positive rate continues to rise when the MBR FDR threshold is increased, as expected.

**Figure 2.**
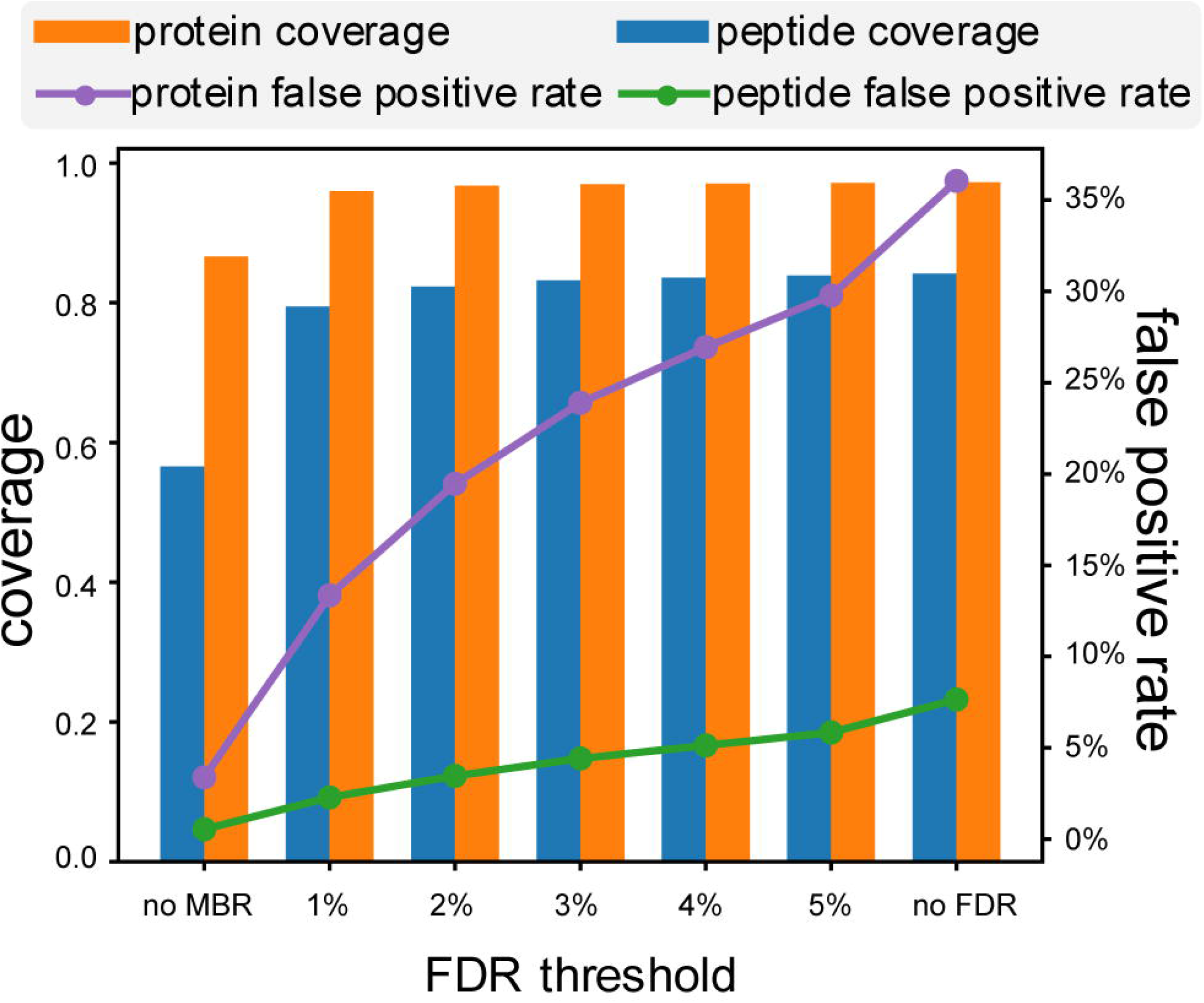
Per-run proteome coverage and observed false positive rate as a function of the model-estimated false discovery rate (FDR) threshold. Coverage is equal to the number of *H. sapiens* peptides/proteins from one run divided by the total number of *H. sapiens* peptide/protein identifications in the entire experiment. The false positive rate is equal to the number of *S. cerevisiae* peptides/proteins from one run divided by the total number of *S. cerevisiae* peptides/proteins.

**Table 2.** Peptides quantified by MaxQuant and IonQuant in analyzing the two-organism dataset with MBR. MSFragger was used to provide identification result for IonQuant. “Sample H” indicates *H. sapiens*-only samples and “Sample HY” indicates samples with a mixture of *H. sapiens* and *S. cerevisiae* proteins. There are 20 runs in each sample type. “MBR+” and “MBR-” indicate that the analysis was performed with and without match-between-runs (MBR), respectively. For each analysis, unique peptide counts (± range of counts) are listed along with per run identification rates (% of all observed peptides found in each run).

| MaxQuant |  |  | IonQuant |  |  |
| --- | --- | --- | --- | --- | --- |
| Total unique *H. sapiens* peptides | 38405 |  | Total unique *H. sapiens* peptides | 45875 |  |
| Sample H, MBR - | 19360±648 | 50.4% | Sample H, MBR - | 26032±499 | 56.8% |
| Sample HY, MBR - | 18945±522 | 49.3% | Sample HY, MBR - | 25683±716 | 56.0% |
| Sample H, MBR + | 31129±637 | 81.0% | Sample H, MBR + | 36450±283 | 79.5% |
| Sample HY, MBR + | 29747±730 | 77.5% | Sample HY, MBR + | 36113±625 | 78.7% |
| Total unique *S. cerevisiae* peptides | 3527 |  | Total unique *S. cerevisiae* peptides | 4610 |  |
| Sample H, MBR - | 20±5 | 0.6% | Sample H, MBR - | 26±6 | 0.6% |
| Sample HY, MBR - | 1848±93 | 52.4% | Sample HY, MBR - | 2597±82 | 56.3% |
| Sample H, MBR + | 98±10 | 2.7% | Sample H, MBR + | 105±16 | 2.3% |
| Sample HY, MBR + | 2858±63 | 81.0% | Sample HY, MBR + | 3625±62 | 78.6% |

In comparing with the results from Skyline, we noticed that using three scores (intensity, retention time difference, and precursor mass error) had a lower false positive rate (**Supporting Table S12**), 5.2% vs 10.4%, than using the default set of scores in training a model using the built-in mProphet. Despite this improvement, mProphet’s false positive rate remained higher than IonQuant’s (2.3%). The peptide numbers in Skyline without MBR are similar to those from IonQuant since both tools were processing the PSMs from MSFragger.

### Improved protein quantification with FDR-controlled MBR

We used four HeLa cell lysate replicates acquired on a timsTOF Pro published by Meier et al. (23) to demonstrate the sensitivity and precision of label-free quantification coupled to FDR-controlled MBR (see **Experimental Procedures**). We previously (22) performed a similar analysis of the same dataset, but without MBR and with protein abundances calculated from peptide ion intensities using top-N peptide approach. In this work we use a new protein abundance calculation module in IonQuant implemented according to the MaxLFQ (15) algorithm (see **Experimental Procedures**).

Table 3 lists the numbers of proteins quantified in at least two runs and the CV from each method. Detailed ion and protein lists can be found in **Supporting Table S2** and **Supporting Table S3.** The results from IonQuant and MaxQuant (both with MaxLFQ method) are shown, which were run under similar settings of requiring either a minimum of 1 or 2 peptide ions in pair-wise ratio calculation in MaxLFQ method (referred to as ‘Min ions’ in IonQuant and ‘LFQ min. ratio count’ in MaxQuant). Enabling MBR (MBR+) improved the number of quantified proteins without a significant increase in protein quantification CV. For example, with min 2 ion setting, IonQuant MBR+ quantified 9% more proteins (5527 vs 5061) while maintaining a CV similar to IonQuant MBR-(3.6% and 3.5%, respectively). Compared to MaxQuant, IonQuant quantified more proteins and with greater precision (lower CVs) in all pair-wise comparisons between the tools under comparable settings. For example, with minimum ion count set to 1, IonQuant with MBR+ quantified 6346 proteins with a CV of 4.0%, compared to 5950 proteins with a CV of 5.3% for MaxQuant with MBR+. IonQuant’s maxLFQ-based protein abundance calculation method also had lower CVs compared to IonQuant with MSstats (47) for peptide to protein intensity roll-up, whereas our initial (top-N peptide based) strategy for protein abundance calculation in IonQuant was inferior to that of MSstats (22) (**Supporting Table S13**).

**Table 3.** Proteins quantified in at least two runs and coefficients of variation (CV) from four HeLa cell lysate replicates. “MBR+” and “MBR-” indicate that the analysis was performed with and without match-between-runs (MBR), respectively..

| Tool |  |  | Proteins quantified | CV |
| --- | --- | --- | --- | --- |
| MaxQuant | MBR- | min 1 ion | 5406 | 5.3% |
|  |  | min 2 ion | 4186 | 4.3% |
|  | MBR+ | min 1 ion | 5950 | 5.3% |
|  |  | min 2 ion | 5073 | 4.7% |
| IonQuant | MBR- | min 1 ion | 5971 | 4.0% |
|  |  | min 2 ions | 5061 | 3.5% |
|  | MBR+ | min 1 ion | 6346 | 4.0% |
|  |  | min 2 ions | 5527 | 3.6% |

We also used the three-organism mixture dataset published by Prianichnikov et al. (24) to demonstrate the accuracy of label-free quantification when FDR-controlled MBR is employed (see **Experimental Procedures**). There are three replicates each of two experimental conditions, where the ratios between the two conditions are 1:1 (*H. sapiens*), 2:1 (*S. cerevisiae*), and 1:4 (*E. coli*). Since these proteomes were mixed at known ratios, we can evaluate the accuracy of the label-free quantification algorithm by comparing the estimated ratio against the ground truth. MaxQuant results published by Prianichnikov et al. (24) were used as a benchmark. We also repeated the analysis with a more recent version of MaxQuant (version 1.6.14.0), a newer reviewed protein database, and parameters as close as possible to those used in MSFragger and IonQuant, and got similar results (**Supporting Figure S1**). We used LFQbench (45) to summarize the analyses and visualize the results (Figure 3 **and Supporting Figure S2**). As expected, both MaxQuant and IonQuant quantified more proteins with MBR than without MBR. IonQuant quantified 6% and 23% more proteins compared to MaxQuant with and without MBR, respectively (Figure 3**, Supporting Table S4, Supporting Table S5**). IonQuant also had fewer outliers than MaxQuant. The peptide level comparison (**Supporting Figure S2**) showed the same trend in comparing IonQuant with MaxQuant.

**Figure 3.**
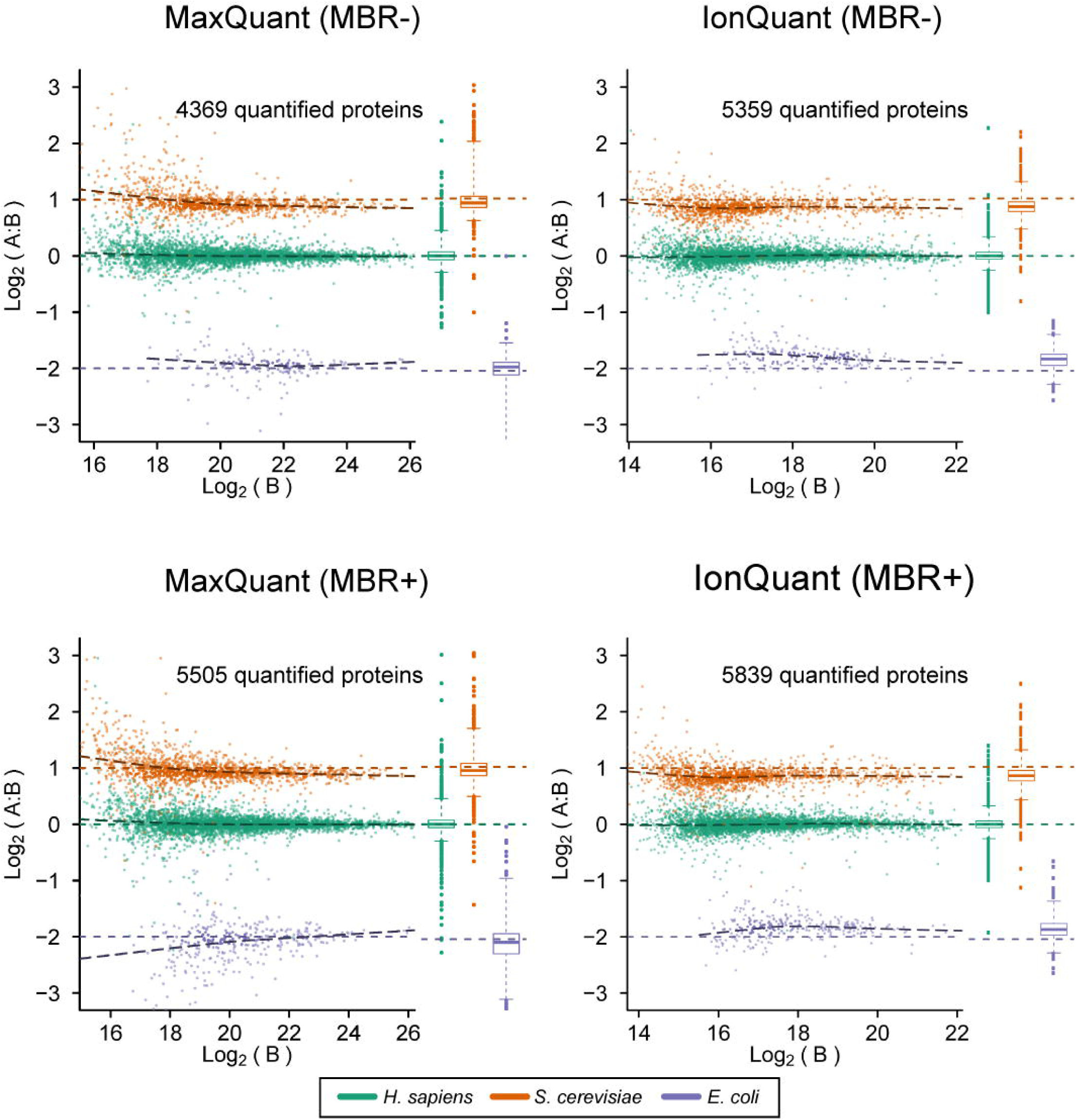
Ground-truth protein quantification results from MaxQuant and IonQuant from a mixture of three different proteomes. MaxQuant results are as published by Prianichnikov et al. 2020. “MBR+” and “MBR-” indicate that the analysis was performed with and without match-between-runs (MBR), respectively. *S. cerevisiae* proteins are shown in orange, *H. sapiens* in green, and *E. coli* in purple. The known ratios of condition A over condition B are 2:1 (*S. cerevisiae*), 1:1 (*H. sapiens*), and 1:4 (*E. coli*). The horizontal colored dashed lines (orange, green, and purple) indicate the true ratios. The black dashed lines are fitted curves from observed ratios. Box plots of the intensities are shown to the right of each scatter plot panel.

### FDR-controlled MBR in single-cell data

We then evaluated the performance of IonQuant with FDR-controlled MBR in single-cell datasets. The first dataset (24) consisted of 5 biological replicates with 1, 3, 10, and 50 cells. In addition, blank runs (0-cells) were also acquired and used as a negative control for MBR. MaxQuant with and without MBR were used as a benchmark.

We first evaluated the number of quantified proteins (proteins with non-zero intensities) (Figure 4(a)). Detailed ion and protein lists can be found from **Supporting Table S6** and **Supporting Table S7**. Of note, MaxQuant with MBR (MBR+) reported on average 68 proteins from a replicate of the blank (0-cell) run, which is much more than MaxQuant MBR-(14 proteins), IonQuant MBR-(19 proteins), and IonQuant MBR+ (31 proteins with 1% FDR). This by itself indicates a noticeable false transfer rate of MaxQuant’s MBR in these data. MSFragger with IonQuant, without MBR (MBR-), identified and quantified a higher number of proteins per sample on average than MaxQuant across all groups of samples. As expected, as the number of cells per sample increases, the average number of proteins quantified per sample, with or without MBR, increases for both MaxQuant and IonQuant. Comparing the numbers from MaxQuant MBR+ and IonQuant MBR+ with FDR set to 1% shows that IonQuant still has a higher number of transferred proteins than MaxQuant, which demonstrates the high sensitivity of IonQuant coupled with MSFragger.

**Figure 4.**
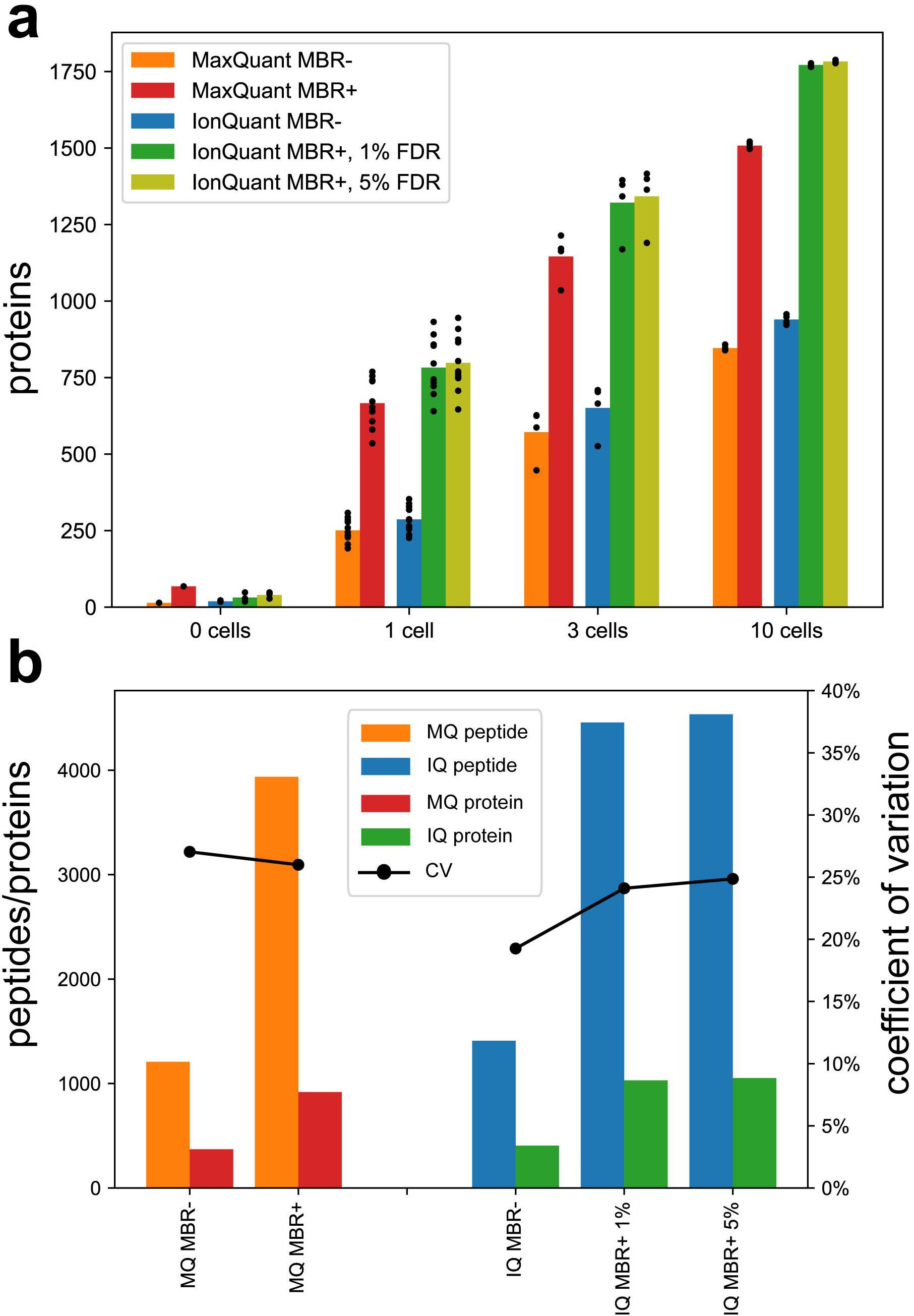
Peptides and proteins from MaxQuant and IonQuant analysis of the single-cell dataset. “MBR+” and “MBR-” indicate that the analysis was performed with and without match-between-runs, respectively. **(a)** Numbers of proteins with non-zero intensities from samples with 0 cells (blank runs), 1 cell, 3 cells, and 10 cells, respectively. Two ion-level MBR false discovery rate (FDR) thresholds (1% and 5%) were applied. Black dots indicate the numbers from individual runs. **(b)** Peptides/proteins quantified in at least two runs and protein quantification coefficient of variation (CV) from 11 replicates of 1 cell samples, as a function of FDR threshold. “MQ” indicates MaxQuant and “IQ” indicates IonQuant. Black curves and dots indicate the CV of the corresponding tool.

Figure 4(b) shows the number of peptides and proteins quantified in at least two runs, and protein quantification CV from analyzing 11 replicates of 1-cell sample with MaxQuant and IonQuant, respectively. Without MBR, IonQuant measured more peptides (1409 vs 1208) and more proteins (406 vs 371), while achieving a lower CV (19.3% vs 27.0%) compared with MaxQuant. With MBR and 1% FDR control, IonQuant also measured more peptides (4457 vs 3937) and more proteins (1030 vs 918) while maintaining a lower CV (24.1% vs 26.0%) compared with MaxQuant.

### FDR-controlled MBR in single-cell data with FAIMS

We used 9 runs (26) from an Orbitrap Eclipse Tribrid mass spectrometer (Thermo Fisher Scientific) coupled with FAIMS to further demonstrate the necessity of controlling FDR for MBR in sparse datasets. There are 3 blank samples containing cell-free supernatant analyzed as negative control, 3 single HeLa cell samples, and 3 samples with 100 HeLa cells to be used as a library for MBR. Each run has two compensation voltages: -55 V and – 70 V. MaxQuant with and without MBR was again used for comparison. Since MaxQuant does not natively support FAIMS data, we split each run into two: one has scans with -55 V and the other has scans with -70 V. In MaxQuant analysis, files with different compensation voltages were assigned to different fractions (i.e., 1 and 3, **Supporting Figure S3**). IonQuant automatically detects and handles FAIMS data, so this manual step is not necessary.

Table 4 shows the number of quantified proteins (proteins with non-zero intensities) from blank and single-cell HeLa samples (the corresponding ions and protein lists can be found in **Supporting Table S8** and **Supporting Table S9).** Both MaxQuant and IonQuant with MBR-identified a relatively large number of proteins in the blank samples (79 and 97 on average per replicate, respectively). This suggests that the blank samples in this experiment cannot be considered as true negative controls for MBR, further highlighting the need for statistical FDR control. While MaxQuant with MBR+ quantified significantly more proteins in the single-cell samples than with MBR-(on average, 1230 vs 557), with MBR+ it also reported on average 492 proteins in the blank samples. In contrast, IonQuant with MBR+ and 1% FDR quantified a comparable number of proteins (on average, 1156) in the single-cell runs as MaxQuant with MBR+, however, the number of quantified proteins in the blank samples has not increased as significantly as with MaxQuant. Applying more lenient MBR FDR thresholds of 2% or 5% in IonQuant results in a significant increase in the number of quantified proteins, while the number of proteins in the blank samples increases as well but still stays below that of MaxQuant with MBR+.

**Table 4.** Number of proteins with non-zero intensities from MaxQuant (MQ) and IonQuant (IQ). “MBR+” and “MBR-” indicate that the analysis was performed with and without match-between-runs (MBR), respectively. The total nonredundant protein count in parentheses, and average proteins per run are outside parentheses.

|  | MQ MBR- | MQ MBR+ | IQ MBR- | IQ MBR+,<br>1% FDR | IQ MBR+,<br>2% FDR | IQ MBR+,<br>5% FDR |
| --- | --- | --- | --- | --- | --- | --- |
| Blank | 79 (152) | 492 (887) | 97 (195) | 153 (314) | 252 (548) | 482 (954) |
| Single-cell<br>HeLa | 557 (853) | 1230 (1902) | 756 (1024) | 1156 (1638) | 1481 (2093) | 2046 (2591) |

Overall, our results above suggest that application of the MBR strategy with no FDR control to sparse datasets, such as single-cell FAIMS data, may result in a high rate of false transfers. IonQuant, with its ability to estimate FDR, provides the users a way to control the rate of false transfers by applying an FDR threshold of their choice. This dataset also invites a discussion regarding a reasonable FDR threshold to apply in different scenarios. In a typical whole cell lysate data, the saturation in the number of quantified proteins is clearly reached at a small FDR threshold (e.g., around 1% FDR in Figure 2(a)). In such datasets, applying a more lenient FDR threshold is likely to reduce the overall quantification accuracy with no noticeable improvement in the number of quantified proteins. Single-cell datasets, on the other hand, are naturally sparser, with more peptides and proteins that can be transferred from other single-cell runs, and especially from the “library” runs (i.e., from boosting samples containing a higher number of cells). In such cases, using a more lenient (e.g., 2%) MBR FDR threshold may be considered, provided that downstream data analysis tools (e.g., for pathway-level analysis) are sufficiently robust toward quantification errors (48).

### Speed of indexing-based MBR in IonQuant

Finally, we compared the computational time required by IonQuant (version 1.5.5) and MaxQuant (version 1.6.14.0), both with MBR enabled. The HeLa dataset (timsTOF Pro) and the two-organism dataset from (Orbitrap Fusion Lumos) were used, comprised of 4 and 40 LC-MS files, respectively (**Experimental Procedures**). For MaxQuant, only jobs related to quantification and MBR were counted (**Supporting Table S10 and Supporting Table S11**). Table 5 displays the run time of these tools in minutes. IonQuant is approximately 19 or 38 times faster than MaxQuant in analyzing the data with or without ion mobility, respectively. The reason that IonQuant exhibits a smaller gain in speed compared with MaxQuant when analyzing the timsTOF Pro data is that most of the IonQuant runtime is spent loading the raw data via the vendor-provided library (22).

**Table 5.** Run time comparison (in minutes) of quantification-related tasks using the HeLa dataset (4 timsTOF Pro runs) and the two-organism dataset (40 Orbitrap Fusion Lumos runs).

|  | HeLa | two-organism |
| --- | --- | --- |
| MaxQuant | 699 | 1056 |
| IonQuant | 37 | 28 |

## Conclusions

Match-between-runs (MBR) is a commonly used approach to quantify additional peptides and proteins by transferring information across different samples. It largely mitigates the missing value problem of DDA-based label-free quantification, increasing data completeness for improved differential analyses. Peptides are transferred from one run to the other by aligning retention time and ion mobility (if applicable). Due to the dynamic range and complexity of proteomic samples, low signal-to-noise ratios and co-isolation interference can result in incorrectly transferred ions. To our knowledge, there was previously no method to control the rate of false transfers in DDA-based MBR in practical settings. To address this issue, we have described a method to estimate and control the FDR for MBR with the help of mixture modeling and the target-decoy concept. We implemented MBR with FDR control in our quantification tool, IonQuant. Our experiments and comparisons with a frequently used tool MaxQuant showed that IonQuant allowed fewer false positive transfers while maintaining high sensitivity. We also highlight the importance of FDR control when MBR is applied to sparse datasets such as those from single-cell FAIMS proteomics experiments. Furthermore, by way of advanced indexing technology, IonQuant performs MBR with unmatched speed, making it well-suited even for analysis of large-scale datasets.

## Abbreviations

LC-MS: liquid chromatography-mass spectrometry
DDA: data-dependent acquisition
DIA: data-independent acquisition
MBR: match-between-runs
FDR: false discovery rate
LDA: linear discriminant analysis
EM: expectation-maximization
LFQ: label-free quantification
CV: coefficient of variation
FAIMS: high-field asymmetric ion mobility spectrometry

## Acknowledgements

This work was funded in part by NIH grants R01-GM-094231 and U24-CA210967. We thank Brett Phinney, Roman Fischer, Tobias Kockmann, and Ying Zhu for useful discussions.

## Data and Software Availability

The two-organism data was published by Lim et al. (18) and can be found at the ProteomeXchange Consortium website (36) with identifier PXD014415. The HeLa cell lysate data was published by Meier et al. (23) and can be found at the ProteomeXchange Consortium website with the identifier PXD010012. The three-organism data was published by Prianichnikov et al. (24) and can be found at the ProteomeXchange Consortium website with identifier PXD014777. The single-cell data was published by Williams et al. (25) and can be found at the ProteomeXchange Consortium website with identifier MSV000085230. MSFragger and IonQuant programs were developed in the cross-platform Java language and can be accessed at http://msfragger.nesvilab.org/ and https://ionquant.nesvilab.org/. Peptide list can be accessed at https://dx.doi.org/10.5281/zenodo.4574598.

## Author Contributions

F.Y. developed IonQuant and its match-between-runs module; F.Y. and A.I.N. analyzed the data; F.Y., S.E.H., and A.I.N. wrote the manuscript with input from all authors; A.I.N. supervised the entire project.

## References

1. Aebersold, R., and Mann, M. (2016) Mass-spectrometric exploration of proteome structure and function. Nature 537, 347–355

2. Aebersold, R., and Mann, M. (2003) Mass spectrometry-based proteomics. Nature 422, 198–207

3. Nesvizhskii, A. I., Vitek, O., and Aebersold, R. (2007) Analysis and validation of proteomic data generated by tandem mass spectrometry. Nat Methods 4, 787–797

4. Nesvizhskii, A. I. (2010) A survey of computational methods and error rate estimation procedures for peptide and protein identification in shotgun proteomics. Journal of proteomics 73, 2092–2123

5. Ludwig, C., Gillet, L., Rosenberger, G., Amon, S., Collins, B. C., and Aebersold, R. (2018) Data-independent acquisition-based SWATH-MS for quantitative proteomics: a tutorial. Molecular systems biology 14, e8126

6. Meier, F., Brunner, A. D., Frank, M., Ha, A., Bludau, I., Voytik, E., Kaspar-Schoenefeld, S., Lubeck, M., Raether, O., Bache, N., Aebersold, R., Collins, B. C., Röst, H. L., and Mann, M. (2020) diaPASEF: parallel accumulation-serial fragmentation combined with data-independent acquisition. Nat Methods 17, 1229–1236

7. Venable, J. D., Dong, M. Q., Wohlschlegel, J., Dillin, A., and Yates, J. R. (2004) Automated approach for quantitative analysis of complex peptide mixtures from tandem mass spectra. Nat Methods 1, 39–45

8. Rosenberger, G., Bludau, I., Schmitt, U., Heusel, M., Hunter, C. L., Liu, Y., MacCoss, M. J., MacLean, B. X., Nesvizhskii, A. I., Pedrioli, P. G. A., Reiter, L., Röst, H. L., Tate, S., Ting, Y. S., Collins, B. C., and Aebersold, R. (2017) Statistical control of peptide and protein error rates in large-scale targeted data-independent acquisition analyses. Nat Methods 14, 921–927

9. Searle, B. C., Pino, L. K., Egertson, J. D., Ting, Y. S., Lawrence, R. T., MacLean, B. X., Villén, J., and MacCoss, M. J. (2018) Chromatogram libraries improve peptide detection and quantification by data independent acquisition mass spectrometry. Nature communications 9, 5128

10. Mueller, L. N., Rinner, O., Schmidt, A., Letarte, S., Bodenmiller, B., Brusniak, M. Y., Vitek, O., Aebersold, R., and Müller, M. (2007) SuperHirn -a novel tool for high resolution LC-MS-based peptide/protein profiling. Proteomics 7, 3470–3480

11. Tsou, C. C., Tsai, C. F., Tsui, Y. H., Sudhir, P. R., Wang, Y. T., Chen, Y. J., Chen, J. Y., Sung, T. Y., and Hsu, W. L. (2010) IDEAL-Q, an automated tool for label-free quantitation analysis using an efficient peptide alignment approach and spectral data validation. Molecular & cellular proteomics : MCP 9, 131–144

12. Zimmer, J. S., Monroe, M. E., Qian, W. J., and Smith, R. D. (2006) Advances in proteomics data analysis and display using an accurate mass and time tag approach. Mass spectrometry reviews 25, 450–482

13. Andreev, V. P., Li, L., Cao, L., Gu, Y., Rejtar, T., Wu, S. L., and Karger, B. L. (2007) A new algorithm using cross-assignment for label-free quantitation with LC-LTQ-FT MS. Journal of proteome research 6, 2186–2194

14. Tyanova, S., Temu, T., and Cox, J. (2016) The MaxQuant computational platform for mass spectrometry-based shotgun proteomics. Nat Protoc 11, 2301–2319

15. Cox, J., Hein, M. Y., Luber, C. A., Paron, I., Nagaraj, N., and Mann, M. (2014) Accurate proteome-wide label-free quantification by delayed normalization and maximal peptide ratio extraction, termed MaxLFQ. Molecular & cellular proteomics : MCP 13, 2513–2526

16. Rieckmann, J. C., Geiger, R., Hornburg, D., Wolf, T., Kveler, K., Jarrossay, D., Sallusto, F., Shen-Orr, S. S., Lanzavecchia, A., Mann, M., and Meissner, F. (2017) Social network architecture of human immune cells unveiled by quantitative proteomics. Nat Immunol 18, 583–593

17. Deshmukh, A. S., Murgia, M., Nagaraj, N., Treebak, J. T., Cox, J., and Mann, M. (2015) Deep proteomics of mouse skeletal muscle enables quantitation of protein isoforms, metabolic pathways, and transcription factors. Molecular & cellular proteomics : MCP 14, 841–853

18. Lim, M. Y., Paulo, J. A., and Gygi, S. P. (2019) Evaluating False Transfer Rates from the Match-between-Runs Algorithm with a Two-Proteome Model. Journal of proteome research 18, 4020–4026

19. Choi, H., and Nesvizhskii, A. I. (2008) Semisupervised model-based validation of peptide identifications in mass spectrometry-based proteomics. Journal of proteome research 7, 254–265

20. Tsou, C. C., Tsai, C. F., Teo, G. C., Chen, Y. J., and Nesvizhskii, A. I. (2016) Untargeted, spectral library-free analysis of data-independent acquisition proteomics data generated using Orbitrap mass spectrometers. Proteomics 16, 2257–2271

21. Tsou, C. C., Avtonomov, D., Larsen, B., Tucholska, M., Choi, H., Gingras, A. C., and Nesvizhskii, A. I. (2015) DIA-Umpire: comprehensive computational framework for data-independent acquisition proteomics. Nat Methods 12, 258–264, 257 p following 264

22. Yu, F., Haynes, S. E., Teo, G. C., Avtonomov, D. M., Polasky, D. A., and Nesvizhskii, A. I. (2020) Fast quantitative analysis of timsTOF PASEF data with MSFragger and IonQuant. Molecular & Cellular Proteomics 19, 1575–1585

23. Meier, F., Brunner, A. D., Koch, S., Koch, H., Lubeck, M., Krause, M., Goedecke, N., Decker, J., Kosinski, T., Park, M. A., Bache, N., Hoerning, O., Cox, J., Rather, O., and Mann, M. (2018) Online Parallel Accumulation-Serial Fragmentation (PASEF) with a Novel Trapped Ion Mobility Mass Spectrometer. Molecular & cellular proteomics : MCP 17, 2534–2545

24. Prianichnikov, N., Koch, H., Koch, S., Lubeck, M., Heilig, R., Brehmer, S., Fischer, R., and Cox, J. (2020) MaxQuant software for ion mobility enhanced shotgun proteomics. Mol Cell Proteomics 19, 1058–1069

25. Williams, S. M., Liyu, A. V., Tsai, C. F., Moore, R. J., Orton, D. J., Chrisler, W. B., Gaffrey, M. J., Liu, T., Smith, R. D., Kelly, R. T., Pasa-Tolic, L., and Zhu, Y. (2020) Automated Coupling of Nanodroplet Sample Preparation with Liquid Chromatography-Mass Spectrometry for High-Throughput Single-Cell Proteomics. Anal Chem 92, 10588–10596

26. Cong, Y., Motamedchaboki, K., Misal, S., Liang, Y., Guise, A., Truong, T., Huguet, R., Plowey, E. D., Zhu, Y., and Lopez-Ferrer, D. (2021) Ultrasensitive single-cell proteomics workflow identifies> 1000 protein groups per mammalian cell. Chemical Science 12, 1001–1006

27. Savitzky, A., and Golay, M. J. (1964) Smoothing and differentiation of data by simplified least squares procedures. Analytical chemistry 36, 1627–1639

28. Freksa, C., Newcombe, N. S., Gärdenfors, P., and Wölfl, S. (2008) Spatial Cognition VI. Learning, Reasoning, and Talking about Space: International Conference Spatial Cognition 2008, Freiburg, Germany, September 15-19, 2008. Proceedings, Springer

29. Millikin, R. J., Solntsev, S. K., Shortreed, M. R., and Smith, L. M. (2018) Ultrafast Peptide Label-Free Quantification with FlashLFQ. Journal of proteome research 17, 386–391

30. Yu, F., Teo, G. C., Kong, A. T., Haynes, S. E., Avtonomov, D. M., Geiszler, D. J., and Nesvizhskii, A. I. (2020) Identification of modified peptides using localization-aware open search. Nature communications 11, 4065

31. Stanley, J. R., Adkins, J. N., Slysz, G. W., Monroe, M. E., Purvine, S. O., Karpievitch, Y. V., Anderson, G. A., Smith, R. D., and Dabney, A. R. (2011) A Statistical Method for Assessing Peptide Identification Confidence in Accurate Mass and Time Tag Proteomics. Analytical Chemistry 83, 6135–6140

32. The, M., and Käll, L. (2020) Focus on the spectra that matter by clustering of quantification data in shotgun proteomics. Nature communications 11, 3234

33. Petyuk, V. A., Qian, W. J., Chin, M. H., Wang, H., Livesay, E. A., Monroe, M. E., Adkins, J. N., Jaitly, N., Anderson, D. J., Camp, D. G., 2nd, Smith, D. J., and Smith, R. D. (2007) Spatial mapping of protein abundances in the mouse brain by voxelation integrated with high-throughput liquid chromatography-mass spectrometry. Genome research 17, 328–336

34. Breen, E. J., Hopwood, F. G., Williams, K. L., and Wilkins, M. R. (2000) Automatic poisson peak harvesting for high throughput protein identification. Electrophoresis 21, 2243–2251

35. Ma, K., Vitek, O., and Nesvizhskii, A. I. (2012) A statistical model-building perspective to identification of MS/MS spectra with PeptideProphet. BMC Bioinformatics 13, 1–17

36. Vizcaino, J. A., Deutsch, E. W., Wang, R., Csordas, A., Reisinger, F., Rios, D., Dianes, J. A., Sun, Z., Farrah, T., Bandeira, N., Binz, P. A., Xenarios, I., Eisenacher, M., Mayer, G., Gatto, L., Campos, A., Chalkley, R. J., Kraus, H. J., Albar, J. P., Martinez-Bartolome, S., Apweiler, R., Omenn, G. S., Martens, L., Jones, A. R., and Hermjakob, H. (2014) ProteomeXchange provides globally coordinated proteomics data submission and dissemination. Nat Biotechnol 32, 223–226

37. Kong, A. T., Leprevost, F. V., Avtonomov, D. M., Mellacheruvu, D., and Nesvizhskii, A. I. (2017) MSFragger: ultrafast and comprehensive peptide identification in mass spectrometry-based proteomics. Nat Methods 14, 513–520

38. Leprevost, F. V., Haynes, S. E., Avtonomov, D. M., Chang, H.-Y., Shanmugam, A. K., Mellacheruvu, D., Kong, A. T., and Nesvizhskii, A. I. (2020) Philosopher: a versatile toolkit for shotgun proteomics data analysis. Nature Methods 17, 869–870

39. Cox, J., and Mann, M. (2008) MaxQuant enables high peptide identification rates, individualized ppb-range mass accuracies and proteome-wide protein quantification. Nature Biotechnology 26, 1367–1372

40. MacLean, B., Tomazela, D. M., Shulman, N., Chambers, M., Finney, G. L., Frewen, B., Kern, R., Tabb, D. L., Liebler, D. C., and MacCoss, M. J. (2010) Skyline: an open source document editor for creating and analyzing targeted proteomics experiments. Bioinformatics 26, 966–968

41. Consortium, U. (2018) UniProt: a worldwide hub of protein knowledge. Nucleic Acids Research 47, D506–D515

42. Keller, A., Nesvizhskii, A. I., Kolker, E., and Aebersold, R. (2002) Empirical statistical model to estimate the accuracy of peptide identifications made by MS/MS and database search. Anal Chem 74, 5383–5392

43. Nesvizhskii, A. I., Keller, A., Kolker, E., and Aebersold, R. (2003) A statistical model for identifying proteins by tandem mass spectrometry. Anal Chem 75, 4646–4658

44. Reiter, L., Rinner, O., Picotti, P., Hüttenhain, R., Beck, M., Brusniak, M. Y., Hengartner, M. O., and Aebersold, R. (2011) mProphet: automated data processing and statistical validation for large-scale SRM experiments. Nat Methods 8, 430–435

45. Navarro, P., Kuharev, J., Gillet, L. C., Bernhardt, O. M., MacLean, B., Rost, H. L., Tate, S. A., Tsou, C. C., Reiter, L., Distler, U., Rosenberger, G., Perez-Riverol, Y., Nesvizhskii, A. I., Aebersold, R., and Tenzer, S. (2016) A multicenter study benchmarks software tools for label-free proteome quantification. Nat Biotechnol 34, 1130–1136

46. Hebert, A. S., Prasad, S., Belford, M. W., Bailey, D. J., McAlister, G. C., Abbatiello, S. E., Huguet, R., Wouters, E. R., Dunyach, J. J., Brademan, D. R., Westphall, M. S., and Coon, J. J. (2018) Comprehensive Single-Shot Proteomics with FAIMS on a Hybrid Orbitrap Mass Spectrometer. Anal Chem 90, 9529–9537

47. Choi, M., Chang, C. Y., Clough, T., Broudy, D., Killeen, T., MacLean, B., and Vitek, O. (2014) MSstats: an R package for statistical analysis of quantitative mass spectrometry-based proteomic experiments. Bioinformatics 30, 2524–2526

48. Paczkowska, M., Barenboim, J., Sintupisut, N., Fox, N. S., Zhu, H., Abd-Rabbo, D., Mee, M. W., Boutros, P. C., Abascal, F., Amin, S. B., Bader, G. D., Beroukhim, R., Bertl, J., Boroevich, K. A., Brunak, S., Campbell, P. J., Carlevaro-Fita, J., Chakravarty, D., Chan, C. W. Y., Chen, K., Choi, J. K., Deu-Pons, J., Dhingra, P., Diamanti, K., Feuerbach, L., Fink, J. L., Fonseca, N. A., Frigola, J., Gambacorti-Passerini, C., Garsed, D. W., Gerstein, M., Getz, G., Gonzalez-Perez, A., Guo, Q., Gut, I. G., Haan, D., Hamilton, M. P., Haradhvala, N. J., Harmanci, A. O., Helmy, M., Herrmann, C., Hess, J. M., Hobolth, A., Hodzic, E., Hong, C., Hornshøj, H., Isaev, K., Izarzugaza, J. M. G., Johnson, R., Johnson, T. A., Juul, M., Juul, R. I., Kahles, A., Kahraman, A., Kellis, M., Khurana, E., Kim, J., Kim, J. K., Kim, Y., Komorowski, J., Korbel, J. O., Kumar, S., Lanzós, A., Lawrence, M. S., Lee, D., Lehmann, K.-V., Li, S., Li, X., Lin, Z., Liu, E. M., Lochovsky, L., Lou, S., Madsen, T., Marchal, K., Martincorena, I., Martinez-Fundichely, A., Maruvka, Y. E., McGillivray, P. D., Meyerson, W., Muiños, F., Mularoni, L., Nakagawa, H., Nielsen, M. M., Park, K., Park, K., Pedersen, J. S., Pich, O., Pons, T., Pulido-Tamayo, S., Raphael, B. J., Reyes-Salazar, I., Reyna, M. A., Rheinbay, E., Rubin, M. A., Rubio-Perez, C., Sabarinathan, R., Sahinalp, S. C., Saksena, G., Salichos, L., Sander, C., Schumacher, S. E., Shackleton, M., Shapira, O., Shen, C., Shrestha, R., Shuai, S., Sidiropoulos, N., Sieverling, L., Sinnott-Armstrong, N., Stein, L. D., Stuart, J. M., Tamborero, D., Tiao, G., Tsunoda, T., Umer, H. M., Uusküla-Reimand, L., Valencia, A., Vazquez, M., Verbeke, L. P. C., Wadelius, C., Wadi, L., Wang, J., Warrell, J., Waszak, S. M., Weischenfeldt, J., Wheeler, D. A., Wu, G., Yu, J., Zhang, J., Zhang, X., Zhang, Y., Zhao, Z., Zou, L., von Mering, C., Reimand, J., Drivers, P., Functional Interpretation Working, G., and Consortium, P. (2020) Integrative pathway enrichment analysis of multivariate omics data. Nature communications 11, 735

