## Supplementary figures for "IonQuant enables accurate and sensitive label-free quantification with FDR-controlled match-between-runs" (from another version of this preprint: version 1, 2020-11-04)

**Figure S1.** Ground-truth protein quantification results from MaxQuant (version 1.6.14.0) with parameters similar to those used in MSFragger and IonQuant. “MBR-” and “MBR+” indicate the analysis was performed without and with match-between-runs (MBR), respectively. *S. cerevisiae* proteins are shown in orange, *H. sapiens* in green, and *E. coli* in purple. The true ratios of three organisms are 2:1 (*S. cerevisiae*), 1:1 (*H. sapiens*), and (1:4) *E. coli*. Box plots of the intensities are shown to the right of each scatter plot panel.

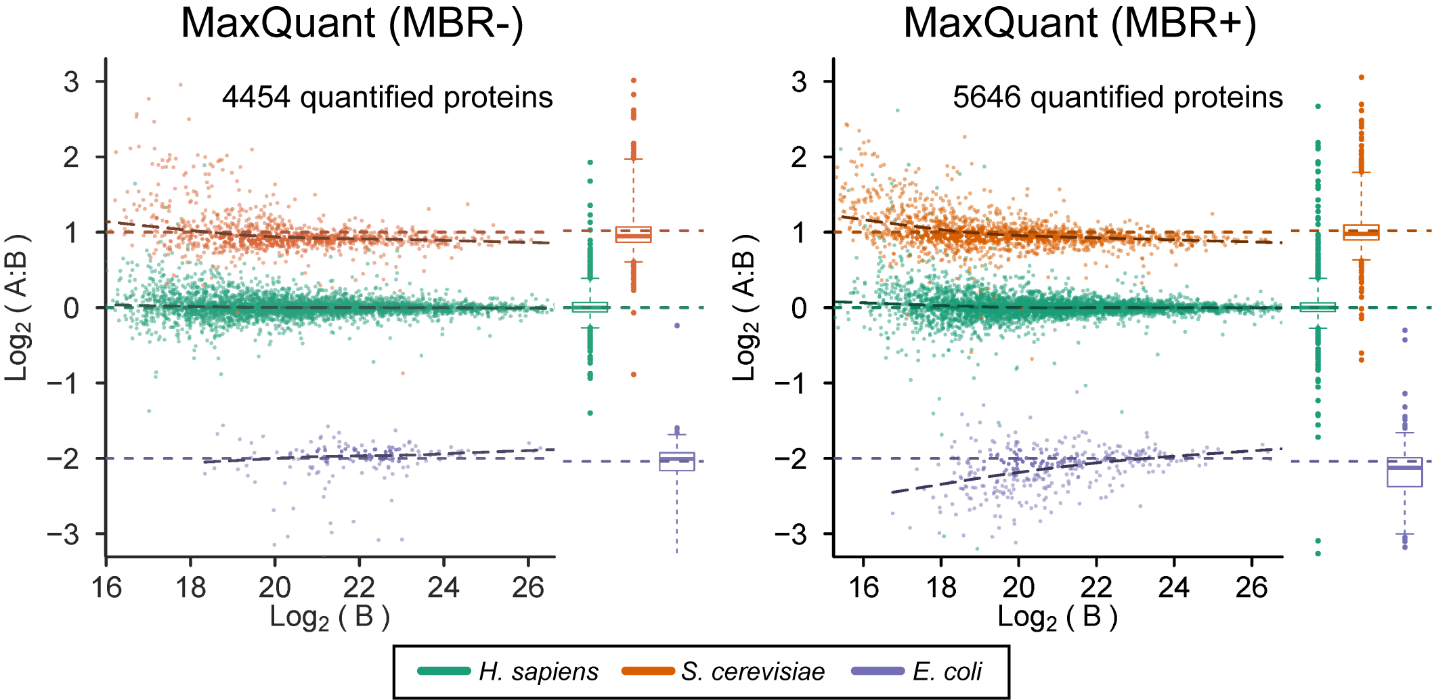

**Figure S2.** Histograms of the target and decoy transferred ions and the fitted distributions. Red indicates targets ions and blue indicates decoys. The yellow lines are fitted target distributions and the green lines are fitted decoy distributions. The first plot is from an LC-MS run published by Meier el al. 2018, the second plot is from a run published by Williams et al. 2020, and the third plot is from a run published by Lim et al. 2019.

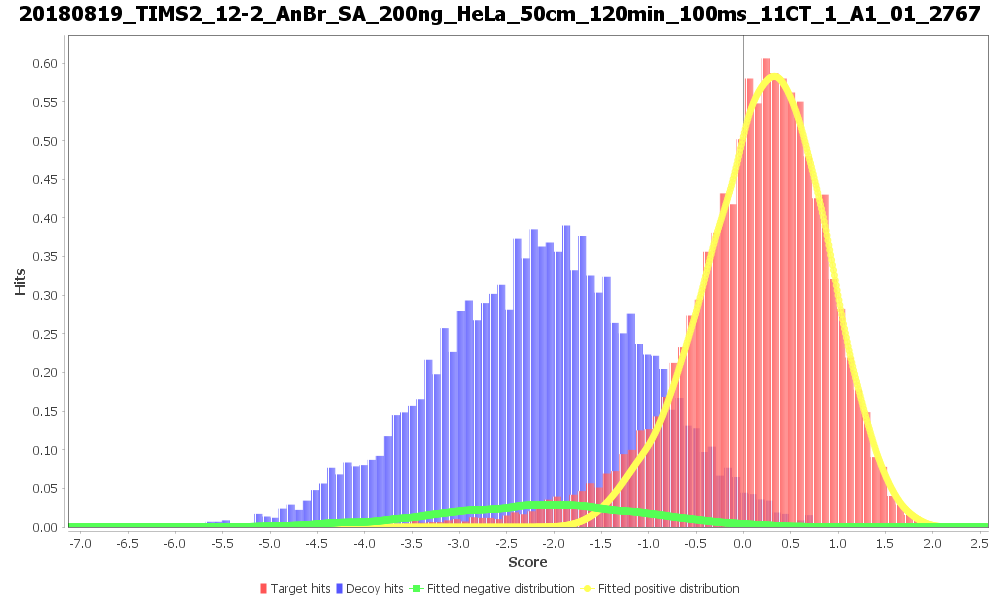

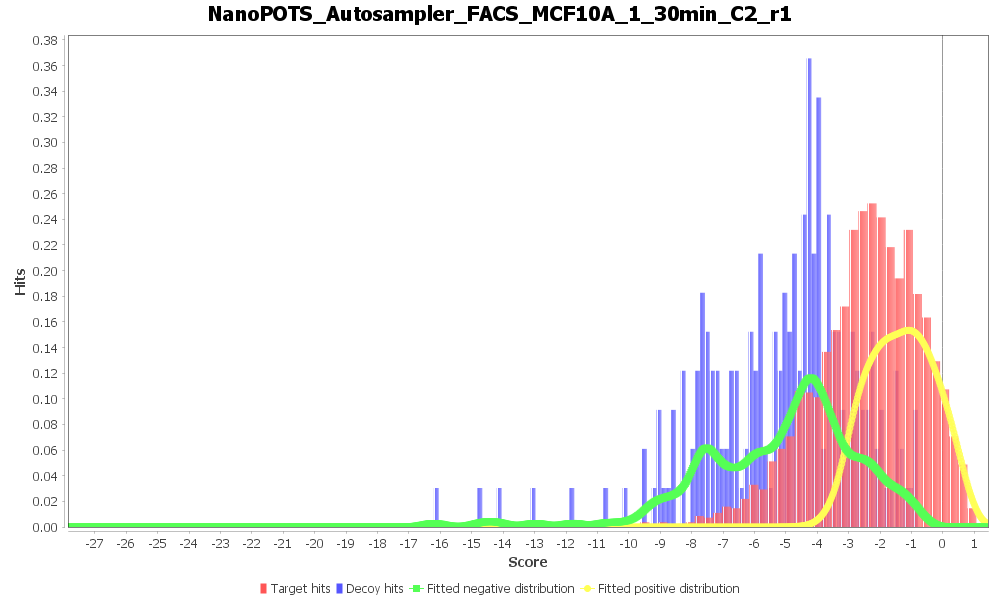

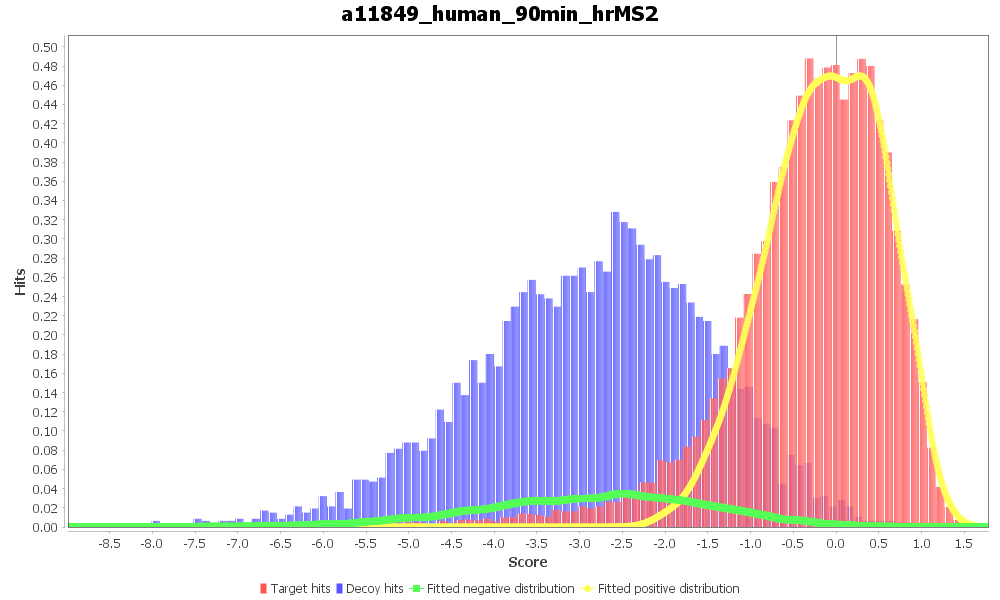
